# Sexual selection on a female copulatory device in an insect with nuptial gifts

**DOI:** 10.1101/2022.03.26.485942

**Authors:** Jessica H. Browne, Darryl T. Gwynne

**Affiliations:** Department of Ecology and Evolutionary Biology, University of Toronto Mississauga, Mississauga, Ontario L5L 1C6, Canada; Department of Biology Mount Allison University, 62 York St. Sackville, NB, E4L 1E2, Canada

**Keywords:** female ornamentation, genital evolution, sexual selection, paternal investments, nuptial gifts, microsatellite analysis

## Abstract

Male genitalia are rapidly evolving structures, often driven sexual selection to increase fertilization success. Although sexual selection on females can be strong in systems where males provide offspring care or feed their mates, sometimes resulting in the evolution of female ornamentation, there are no actual estimates of direct sexual selection on female genitalia. In a New Zealand ground weta, *Hemiandrus pallitarsis* (Orthoptera: Ensifera, Anostostomatidae), females possess a genitalic device (the accessory organ) that is necessary for successful copulation and the acquisition of glandular food-gifts from males. These nutritious gifts are known to result in sexual competition among females in other ensiferan species. In ground weta, the gifts are probably important in avoiding starvation during a months-long period when caring for (their lifetime production of) eggs and offspring. Here, we test the hypothesis that the accessory organ is a sexually selected device in *H. pallitarsis* by measuring the female Bateman gradient and directional sexual selection on the accessory organ. Using newly developed and characterized microsatellite loci, we analyze offspring and/or stored sperm to estimate female mating frequency for the first time in ground weta. As predicted, we found positive Bateman gradients for females, and some evidence of directional sexual selection on accessory organ length. Although organ length does not correlate well with female fecundity, it may increase mating success by indicating her condition and thus quality of her offspring care.

## Introduction

Genitalia and associated structures (secondary genitalia) are some of the most rapidly evolving animal traits, often exhibiting extreme variation among closely related species (Eberhard 1985). While genital diversity has been suggested to function in species isolation (“lock and key” hypothesis; Mayr 1963; Eberhard 1985), genital evolution in males is thought to be driven mainly by sexual selection for increased fertilization success (Eberhard 1985). Because females are typically more selective when mating (Darwin 1871; Trivers 1972), elaborate male genitalia may have evolved in the context of cryptic female choice, where males with the most “attractive” stimulatory genitalia experience the greatest fertilization success (Lloyd 1979; Thornhill 1983; Eberhard 1985; Simmons 2001). Alternatively, genitalia may function in manipulating female mating behaviour and/or her sperm stores (e.g., grasping devices that coerce copulations or prevent female remating; Arnqvist and Rowe 2005). Although much of the focus has been on male traits, females are not passive in this process (Sloan and Simmons 2019). In response to, for example, male grasping or piercing genitalia, there can be co-evolution of female genitalia to reduce potential costs of mating (e.g., Arnqvist and Rowe 2005; Morrow and Arnqvist 2003) or to bias fertilization success in favour of high-quality males (Eberhard 1985; Sloan and Simmons 2019).

In contrast to these examples of “typical” mating roles, the factors influencing male and female fitness are expected to differ when males make valuable investments in paternal care or nuptial gifts. Females receiving these direct benefits are expected to seek additional matings, while the potential for frequent mating by males can be limited by investment cost, resulting in male preferences and/or direct sexual competition among females (Trivers 1972; Williams 1975; Parker and Simmons 1996; Bonduriansky 2001; Gwynne 2016; Herridge et al. 2016). In these systems, especially those where females depend on male investments for offspring production, female competition can cause strong sexual selection on females for traits that are attractive to males (Tobias et al. 2012; Herridge et al. 2016; Hare and Simmons 2018). Such traits are well-known both in certain vertebrates with paternal care (e.g., sex-specific female ornaments in pipefish: Flanagan et al. 2014) and in insects such as empid dance flies (legs, wing, or abdominal ornaments; Cumming 1994; Murray et al 2022) where females rely on male-provided nutrition (insect prey items) to develop their eggs (Downes 1970). Also, in katydids (Orthoptera: Ensifera; Tettigoniidae) where males supply large spermatophore gifts (Vahed and Gilbert 1996), there is sexual selection on females when they rely on these gifts for nutrition, which impose additional mating costs for males (Gwynne 1981; 1984; Simmons and Bailey 1990; Gwynne and Simmons 1990; Simmons 1992; Robson and Gwynne 2010). In *Kawanaphila nartee*, females with larger ears are faster in the race to reach singing males (Gwynne and Bailey 1999) and there is positive selection for ear size, a sexually dimorphic trait in this species, only when food is limited (Hare and Simmons 2021).

Virtually unstudied is direct sexual selection on sex-specific female genitalic traits where copulation is associated with acquiring goods or services from males. As in males, female genital traits may be expected to evolve either in the context of mate choice – such as stimulating males into copulating – or coercing males into supplying more food. Suggested examples are (1) the interspecifically varying ridges on female subgenital plates of *Kawanaphila* katydids (Rentz 1993) that grasp the male’s genitalia, perhaps allowing longer copulation duration and thus the delivery of a larger spermatophore gift (Gwynne 2001); and (2) a penis-like female genital modification in two genera of psocopteran cave insects, *Neotrogla* and *Afrotrogla*. Male psocopterans can transfer nutritious semen to females during copulation, which is thought to be an important source of nutrients (Wearing-Wilde 1996; Yoshizawa et al. 2018). The penis-like gynosome, inserted into the male’s genital opening, anchors him during copulation (Yoshizawa et al. 2014), and is a device likely under sexual selection to allow females to prolong the duration of copulation and thus the amount of food-gift received (Yoshizawa et al. 2014; Yoshizawa et al. 2018).

Our study species has female “secondary” genitalia, abdominal devices found in a clade of New Zealand “short-tailed” ground weta (*Hemiandrus*, Anostostomatidae, Orthoptera: Trewick et al. 2020). As in *Neotroglea* (Yoshizawa et al. 2014), male ground weta transfer nutrients to their mates, in this case, a glandular spermatophylax part of the spermatophore (as in katydids; Gwynne 1995) that the mating male adheres to the female’s abdomen (separate from the sperm capsule) and is eaten by her after copulation (Gwynne 2004; Browne and Gwynne 2022). The female device is an elongated abdominal modification on the 6^th^ sternite, known as an accessory organ (Salmon 1950). In contrast to other species of *Hemiandrus* that, like most ensiferan species, abandon their eggs after ovipositing into the substrate (Gwynne 2004; Taylor Smith et al. 2013), short-tailed females care for their eggs and newly-hatched offspring (their lifetime reproduction) in an underground chamber for a period of 5-6 months (hence the “short-tail” reduced ovipositor; Gwynne 2004; Gwynne 2005; Taylor Smith 2015) and then die. Care-providing females do not eat, so male mating gifts likely represent important nutrition to avoid starvation during the lengthy care period, potentially leading females to compete for mating (feeding) opportunities (Gwynne 2004; Gwynne 2005). Because of this, the accessory organ, positioned close to where the sticky spermatophylax gift is adhered, is expected to be involved in the acquisition of gifts.

Male genitalia in many weta (Anostostomatidae) and some related ensiferans, include genital hooks (falci) on the dorsal side of the 10^th^ segment (Johns 2001) which hook onto the edge of one of the of the female’s sternites during copulation (the male is positioned beneath the female: Gwynne 2002). This is likely an ancestral behaviour, used to grip the female for stability or to coerce her into copulation (e.g., Weissman 2001). While some weta species have sclerotised cuticle on the sternite edge that receives the male hooks, female short-tailed ground weta are unique in possessing the accessory-organ modifications on the 6^th^ sternite, to which the male genitalia attach, but do not hook (D. Gwynne, personal observation). Females of our study species, *H. pallitarsis*, possess the most elaborate accessory organ (fig. 1), which protrudes from the abdomen. After pairing (female mounted above male), males first make genitalic contact with the accessory organ then move their genitalia back to attach to the female’s primary genitalia when they insert the sperm ampulla (Gwynne 2004; Gwynne 2005). Following this, the male detaches from her primary genitalia, but his genitalia remain attached to her abdominal accessory organ while he extends paired phalli that deposit the bilobed spermatophylax gift (Gwynne 2002; Gwynne 2004; Gwynne 2005). During gift transfer, the male’s paired falci press onto the dorsal surface of the accessory organ, while his paired epiprocts appear to be inserted ventral to the organ, and the distal edge of the male’s 8^th^ abdominal segment pushes into the region between the female’s 5^th^ and 6^th^ sternites (photographic evidence: DTG unpublished). The placement of the spermatophylax gift anterior to the accessory organ suggests that it is not a holding device (see Gwynne 2002) for the gift (Gwynne 2005). However, it is necessary for mating (Gwynne 2005) as copulation fails when the female organ is experimentally removed (Gwynne 2005). Thus, this device has not evolved to thwart copulation attempts (as in some gerrid water striders: Arnqvist and Rowe 2005).

**Figure 1.**
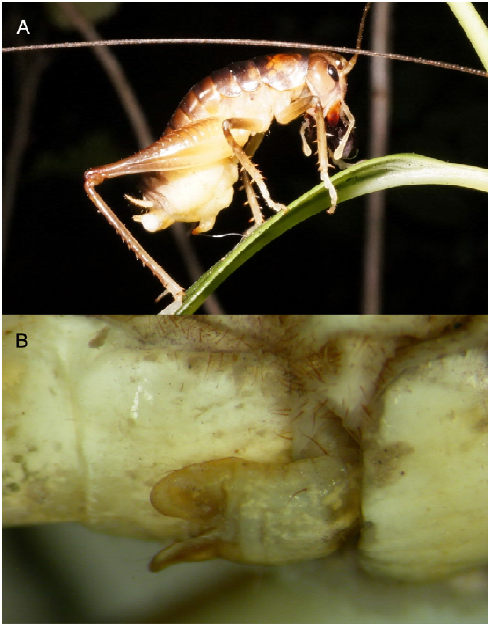
Lateral view of the extended female secondary genitalic accessory organ (6^th^ sternite, located between the hind and middle legs) in A) a newly-mated female *H. pallitarsis* and B) a close-up of the organ. Photos by Darryl Gwynne.

Although the female accessory organ is involved in transfer of the ejaculate (sperm and the gift in ground weta), there is some evidence that it is sexually selected to secure matings as envisioned by Eberhard (1990) for male genitalia (examples: fish, Brooks and Caithness 1995; lizards, Bohme 1983; flies,. 2021; Briceño et al. 2010; and katydids, Wulff and Lehmann 2016). First, extensive interspecific variation in the accessory organs of short-tailed ground weta (Johns 2001; Trewick et al. 2020) reflects the typical pattern in male animals (Gwynne 2005) where external genitalia vary greatly across closely related species. Variation in female accessory organs is probably not related to species isolation (“lock and key” hypothesis; Mayr 1963; Eberhard 1985) since there is greater support for sexual selection on complex genitalia in male insects (Eberhard 1985; Gwynne 2005) and, for ground weta, mating interactions between species are more likely prevented by the species-specific precopulatory drumming behaviour of males and females that precede pairing (Gwynne 2004; Gwynne 2005; Chappell et al. 2012). Furthermore, there is little overlap in ground weta species ranges (Trewick and Bland 2011; Trewick et al. 2020), which makes species mis-pairing unlikely. Also consistent with the sexual selection hypothesis, male ground weta make close contact with the accessory organ prior to mating and have been observed to reject females after probing her mid-abdomen (Gwynne 2005). As females with longer accessory organs tend to have a greater number of eggs that develop to the nymphal stage (female body size controlled; Gwynne 2005), this trait may help females obtain matings (gifts) by signalling quality to males (as in Andersson 1994; Berglund et al. 1997; Barry 2015). Thus, one hypothesis is that this structure evolved via male choice in the context of female mate/gift acquisition (Gwynne 2005),

As the female organ is inserted between two parts of the male genitalia, another possibility is that it has evolved by sexual selection on females to anchor between the male genital falci and epiprocts, and thus manipulate mating or spermatophylax-gift transfer, in a similar manner to the anchoring female penis of *Neotroglea* bark lice suggested by Yoshizawa et al. (2014) as a sexually-selected organ to obtain nutritious ejaculates. Gwynne’s (2005) finding that accessory organ length correlates with the number of eggs that develop to the nymphal stage in *H. pallitarsis* is also consistent with this hypothesis if accessory organs result in a longer copulation and more nuptial food. However, there is no direct evidence that any female copulation devices such as accessory organs or female penes are ornaments subject to sexual selection. Here, we test this by measuring sexual selection on female ground weta *H. pallitarsis*. Using estimates of female mating success from microsatellite analysis of stored sperm and measures of lifetime female reproductive success from females that cared for their broods in the lab, we determine the degree of sexual selection on females (Bateman gradient) and estimate directional sexual selection on several accessory organ traits. If the accessory organ is sexually-selected in the context of gift acquisition, we predict directional selection on accessory organ size (length; Gwynne 2005) and that this trait will correlate with female fecundity. Characteristic of sexual selection on a sex (Bateman 1948; Arnold 1994), we also expect there to be a positive relationship between female mating success and reproductive success (Bateman gradient).

## Methods

### Specimen collection: Bateman gradients

To determine Bateman gradients for female ground weta *(H. pallitarsis*), females were collected in 2002 (n=10) and 2017 (n=17) from two main sites on the North Island of New Zealand: Kiriwhakapapa trail near Masterton (−40.807627, 175.546532) and private gardens in Palmerston North (−40.413909, 175.662814). Collections took place from mid-February to early March, nearing the end of the ground weta mating season (Gwynne 2004). Live specimens were transported back to our lab in Toronto, Canada to oviposit and obtain measures of lifetime reproductive success from caring females. This was a lengthy process that involved placing females in artificial brood chambers formed from potter’s clay (2002) or soil (2017) and allowing 2-3 months for them to lay eggs, as well as an additional 5-6 months for eggs to develop, during which time females were not fed. Several females died during this period, four of which were excluded from further analysis because both her and her eggs decayed rapidly (final n=23). Once eggs began hatching, mothers and their broods were preserved at -20°C to prevent consumption of dead or developing tissue. We counted the total number of eggs laid as well as the number of hatched (1st instar nymphs) or nearly hatched (eggs with visible eye spots) offspring, which we considered “developed eggs”. To obtain the sperm from mates that did not fertilize offspring, we dissected female sperm storage organs, isolated the sperm pellet (using >70% ethanol; Tripet et al. 2001), and stored at -20°C. We then photographed females under a dissecting scope fitted with a digital camera and measured pronotum length as an estimate of body size (as in Taylor Smith 2015).

### Specimen collection: Sexual selection on accessory organs

To measure sexual selection on accessory organs, additional *H. pallitarsis* females (n=58) were collected in 2012, 2016, 2017, and 2019 near the end of the mating period from the same two sites: Kiriwhakapapa and Palmerston North. Because our primary interest here was not reproductive output, these females were frozen immediately following capture and stored in strong ethanol (70-95%). This method allowed us to avoid a time-consuming and risky offspring-rearing period and ensured that both female phenotype and the quality of sperm stores were not compromised. For this same reason, we did not include the 23 females that laid eggs (used for Bateman gradient estimates) in our measures of selection on accessory organs. We photographed preserved females under a dissecting scope and measured pronotum length (body size; Taylor Smith 2015) as well as accessory organ length and width (fig. 2) using ImageJ software (Schneider et al. 2012). Prior to measurement, all accessory organs were removed and soaked in a 5% Decon 90 solution for 1 hour at 40ºC to soften and restore the structures (Banks and Williams 1972), as many were bent and brittle after prolonged storage in ethanol. We then dissected the spermathecae from all females, isolated the sperm contents using strong ethanol (Tripet et al. 2001), and stored them at -20°C. During the dissection, we also counted the number of developing ova in the female’s abdomen.

**Figure 2.**
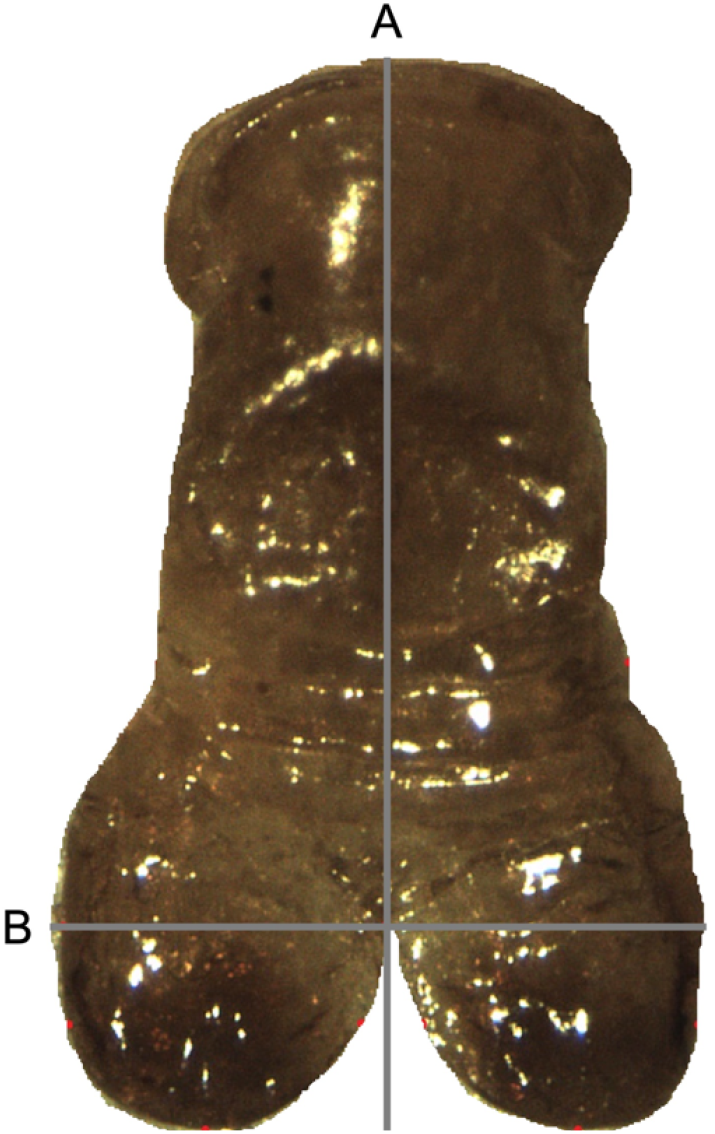
Modified 6^th^ sternite, the accessory organ, on the abdomen of female *H. pallitarsis* ground weta. Image shows the structure removed from the body and the two measurements taken: A) accessory organ length, from the point where the organ attaches to the body to the furthest point on the distal end and B) accessory organ width, left to right on the widest point of the distal end.

### Microsatellite analysis

Prior to conducting genetic analysis, we assessed amplification quality as well the degree of polymorphism for the nine candidate loci in our microsatellite library (developed by Genetic Marker Services, Brighton, UK for *H. pallitarsis*; provided in Online Resource 1: Table S1) We used polymerase chain reaction (PCR) to amplify DNA samples extracted from adult hind leg tissue, including the females in the current study plus an additional 24 males. Using GENEPOP version 4.2 (Raymond and Rousset 1995), we determined that the degree of genetic differentiation (F_st_) was low between our two study sites (populations) and identified three polymorphic loci (>20 alleles) with a low frequency (< 20%; Dakin and Avise 2004) of null alleles (Dempsters EM method; heterozygote deficiency test) and non-specific amplification (wet80, wet 81, and wet89; details in Online Resource 1: Table S2). Targeting these loci, we PCR-amplified samples of stored sperm and offspring. We extracted DNA from stored sperm using the Qiagen DNeasy Blood and Tissue Extraction kit (Qiagen, Hilden, Germany), and added 12μL DTT to each sample to improve lysis of sperm cells. We extracted DNA from offspring tissue (nymph or egg) using a Proteinase K based extraction method with multiple ethanol washes. The resulting concentration of DNA was much higher for offspring relative to sperm and so we diluted offspring samples to roughly 50ng/μL. Each PCR reaction contained 5.75μL sterile filtered water, 1μL 10X PCR reaction buffer, 0.2μL 10mM dNTP mixture, 0.5μL of forward primer (each with fluorescent labels), 0.5μL reverse primer, and 1μL of template DNA. Thermocycling was done using an Eppendorf Mastercycler with the following temperature regime: 2 minutes at 94°C followed by 35 cycles of 30 seconds at 94°C, 30 seconds at 60°C, and 15 seconds at 72°C, then an additional 3 minutes at 72°C. PCR products were sent to The Centre for Applied Genomics at Sick Kids Hospital in Toronto, ON for fragment analysis on an Applied Biosystems 3730xl capillary sequencer. The resulting electropherograms were examined using GeneMarker V1.97, which allowed us to identify fluorescence peaks, representing alleles.

### Estimating female mating success

For measures of the Bateman gradient, we used a combination of paternity data and the remaining stored sperm to estimate female mating frequency. Based on offspring genotypes, we first determined the minimum number of sires for each brood using the parentage program GERUD 2.0 (Jones 2005). Any alleles found in female’s sperm stores that were not present in the offspring genotypes were assumed to be additional mates and were factored into a female’s total mating frequency. We estimated the most probable number of additional mates at each locus using a method described by Bretman and Tregenza (2005). This method (See Bretman and Tregenza 2005; Herridge 2016 thesis for details) makes use of known allele frequencies in the population to determine the probability of observing a particular set of alleles given different numbers of contributing males. This provides a less conservative estimate than simple allele counting, which assumes heterozygosity (two alleles indicate one individual). Because estimates of females mating success incorporated offspring data, we could not remain blind to reproductive success (the response) when collecting these data. One of our specimens was known to have mated, as she was found with remnants of a spermatophylax gift in her mouthparts; however, the male (who remains close by while she consumes the gift) failed to inseminate this female as his genotype did not appear in her sperm stores, nor her offspring. Because acquisition of male-provided materials (in this case the spermatophylax gift) is the key factor affecting female fitness (Clutton-Brock 2007), an additional mate was added to this female’s total mate number for our estimates of the Bateman Gradient.

To measure directional sexual selection on accessory organs, we estimated female mating frequency from stored sperm alone. Using all male alleles detected in stored sperm, we determined the most probable mate number (see Bretman and Tregenza 2005) obtained by each female. Unfortunately, due to poor amplification at all loci, we were unable to estimate the mate number for four females. In cases where an allele was found in sperm stores that was not previously detected in the population, it was assumed to be rare and assigned a frequency equal to the least common allele known in that population. We were blind to estimates of female mating frequency as they were made without knowledge of phenotype.

### Statistical analysis: Bateman gradients

We measured the Bateman gradient for females using our estimates of mating frequency from stored sperm and the number of offspring in single broods that each female cared for before dying (lifetime reproductive success). We calculated standardized Bateman gradients (β′_ss_; Bateman 1948; Arnold 1994) using an ordinary least squares (OLS) regression in which relative number of mates (mean-standardized) was the predictor variable and relative reproductive success (mean-standardized) was the response variable. A separate regression model was run for each of our measures of reproductive success: the number of eggs laid and the number of eggs that developed to the nymphal stage (determined by hatching or visible eye spots). We measured pronotum length as an indicator of body size (e.g., Taylor Smith 2015), however decomposition of several females in their brooding chambers prevented us from obtaining data for all specimens. Using the available pronotum data (n=20), we included body size in our regression model to determine whether this explained significant variation in reproductive success or altered measures of the Bateman gradients Due to our low sample size, and deviations from normality (assessed visually with Q-Q plots), we used permutation tests to obtain p-values. We ran 5000 permutations to generate a null distribution of possible Bateman gradients (regression coefficients) from our data and compared this to our observed values to determine if they were significantly different from those expected by chance. Due to apparent variation between our collection sites and years, we assessed whether measures of the Bateman gradient interacted with either of these variables. Using separate regression models for each variable, we included site and year as interaction terms and used p values derived from multiple least squares regression to assess significance. We did not use permutation tests to obtain p values for interaction terms, as this approach does not preserve important associations in the data, leading to inflated error rates (Bužková et al. 2011; Bužková 2016; Foster et al. 2016).

### Statistical analysis: Sexual selection on female accessory organs

We measured sexual selection acting directly on female accessory organs using female phenotype (body size, accessory organ length, accessory organ width) and estimates of mating frequency from stored sperm. We obtained linear selection gradients (indicative of directional selection) using multiple least squares regression (MLS) in which trait values (z-score standardized) were predictor variables and relative (mean-standardized) number of mates was the fitness response variable. Low sample size limited the complexity of our regression model; thus, we were unable to include quadratic and correlational terms (Lande and Arnold 1983) in our analysis. To determine whether observed selection gradients were statistically significant, we again used permutation tests with 5000 randomizations (as in Kelly & Gwynne 2023). Due to apparent variation in the effect of female body size as well as accessory organ width between our two collection sites, we tested for interactions using MLS regression. We ran individual models for each female trait, in which the interaction with site was included as a predictor, and used this to derive both effect sizes and p-values for interaction terms (see Bužková et al. 2011; Bužková 2016; Foster et al. 2016 for limitations of permutation tests)

Finally, in order to assess whether the accessory organ has the potential to signal female fecundity, we ran another MLS regression model with z-score standardized trait values (body size, accessory organ length, accessory organ width), but used relative (mean-standardized) number of developing eggs in the abdomen (not yet laid) as the response variable. Again, we used permutation tests with 5000 randomizations to obtain p-values and tested for interactions between site and either body size or accessory organ width (due to visual variation) using MLS regression.

## Results

### Bateman gradients

Consistent with our prediction that female ground weta are subject to sexual selection, we found evidence of significant positive Bateman gradients in one of our two collection years (table 1). Although Bateman gradients using pooled data were found to be positive, they were not statistically significant (table 1) and showed considerable variation across our two collection sites and years, especially when using the total eggs laid (fig. 3). While there were no significant interactions when using the number of developed eggs, there was a significant interaction with year when using eggs laid (table 1), leading us to analyze data separately for each year. The Bateman gradient was high in the year 2017 and appears to drive the overall positive trend found between mating success and the number of eggs laid (table 1; fig. 3).

**Table 1.**
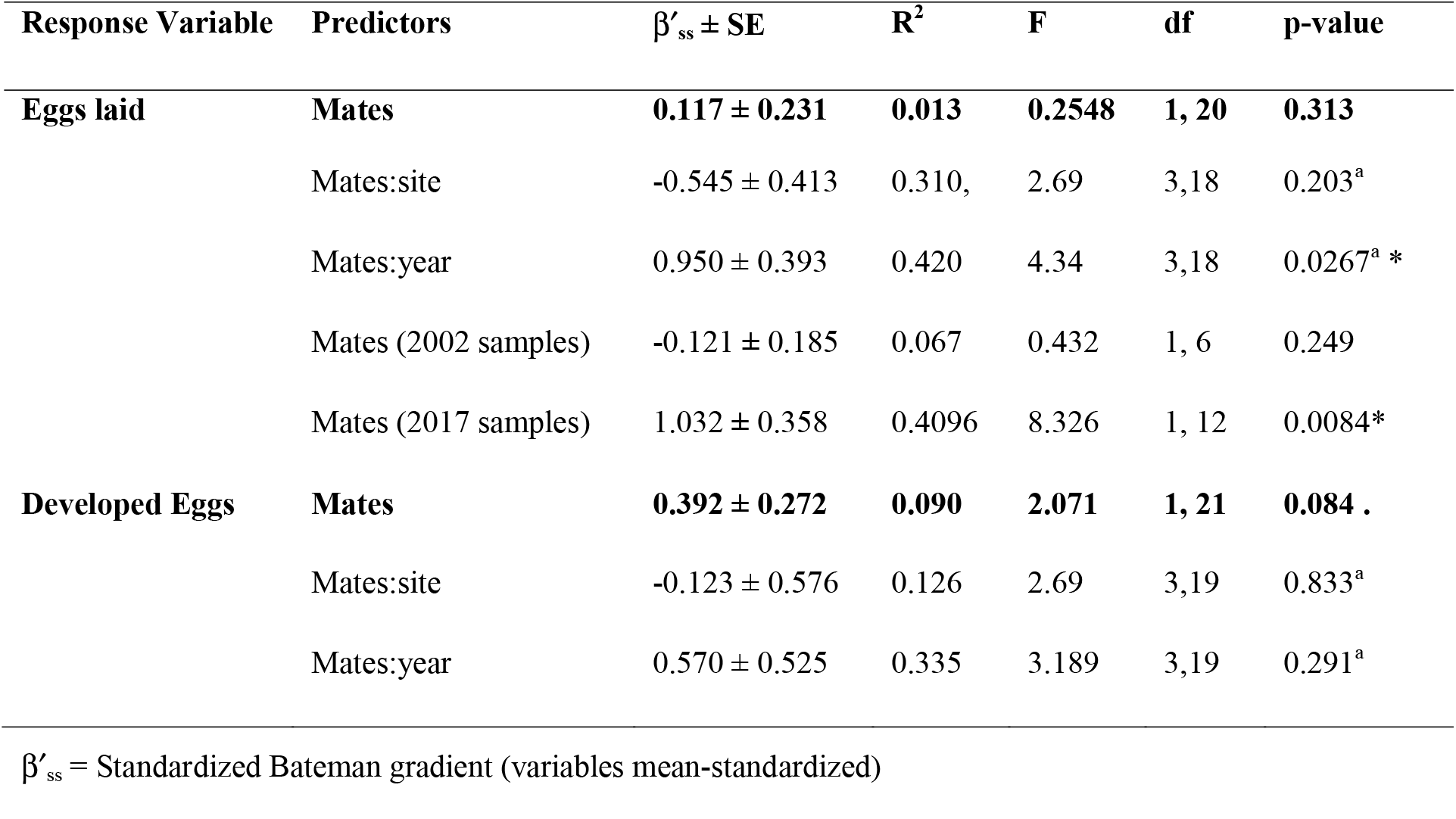
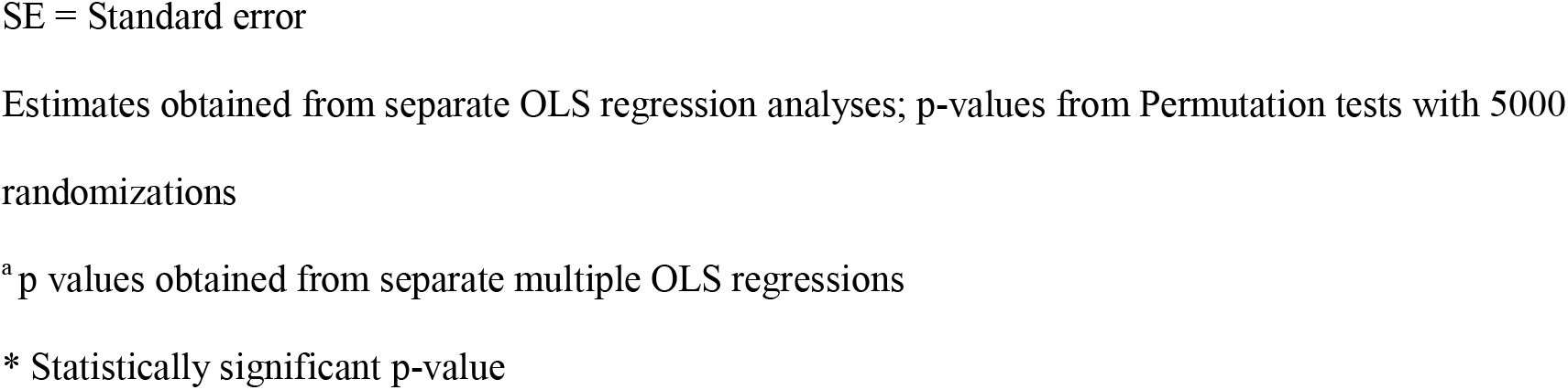
Standardized Bateman Gradients (β′_ss_) for wild-caught female ground weta, *H. pallitarsi*s.

| Response Variable | Predictors | $\beta'_{ss} \pm SE$ | $R^2$ | F | df | p-value |
| --- | --- | --- | --- | --- | --- | --- |
| **Eggs laid** | **Mates** | **0.117 ± 0.231** | **0.013** | **0.2548** | **1, 20** | **0.313** |
| | Mates:site | -0.545 ± 0.413 | 0.310, | 2.69 | 3,18 | 0.203<sup>a</sup> |
| | Mates:year | 0.950 ± 0.393 | 0.420 | 4.34 | 3,18 | 0.0267<sup>a</sup> * |
| | Mates (2002 samples) | -0.121 ± 0.185 | 0.067 | 0.432 | 1, 6 | 0.249 |
| | Mates (2017 samples) | 1.032 ± 0.358 | 0.4096 | 8.326 | 1, 12 | 0.0084* |
| **Developed Eggs** | **Mates** | **0.392 ± 0.272** | **0.090** | **2.071** | **1, 21** | **0.084 .** |
| | Mates:site | -0.123 ± 0.576 | 0.126 | 2.69 | 3,19 | 0.833<sup>a</sup> |
| | Mates:year | 0.570 ± 0.525 | 0.335 | 3.189 | 3,19 | 0.291<sup>a</sup> |

$\beta'_{ss}$ = Standardized Bateman gradient (variables mean-standardized)
SE = Standard error
Estimates obtained from separate OLS regression analyses; p-values from Permutation tests with 5000 randomizations
<sup>a</sup> p values obtained from separate multiple OLS regressions
\* Statistically significant p-value

**Figure 3.**
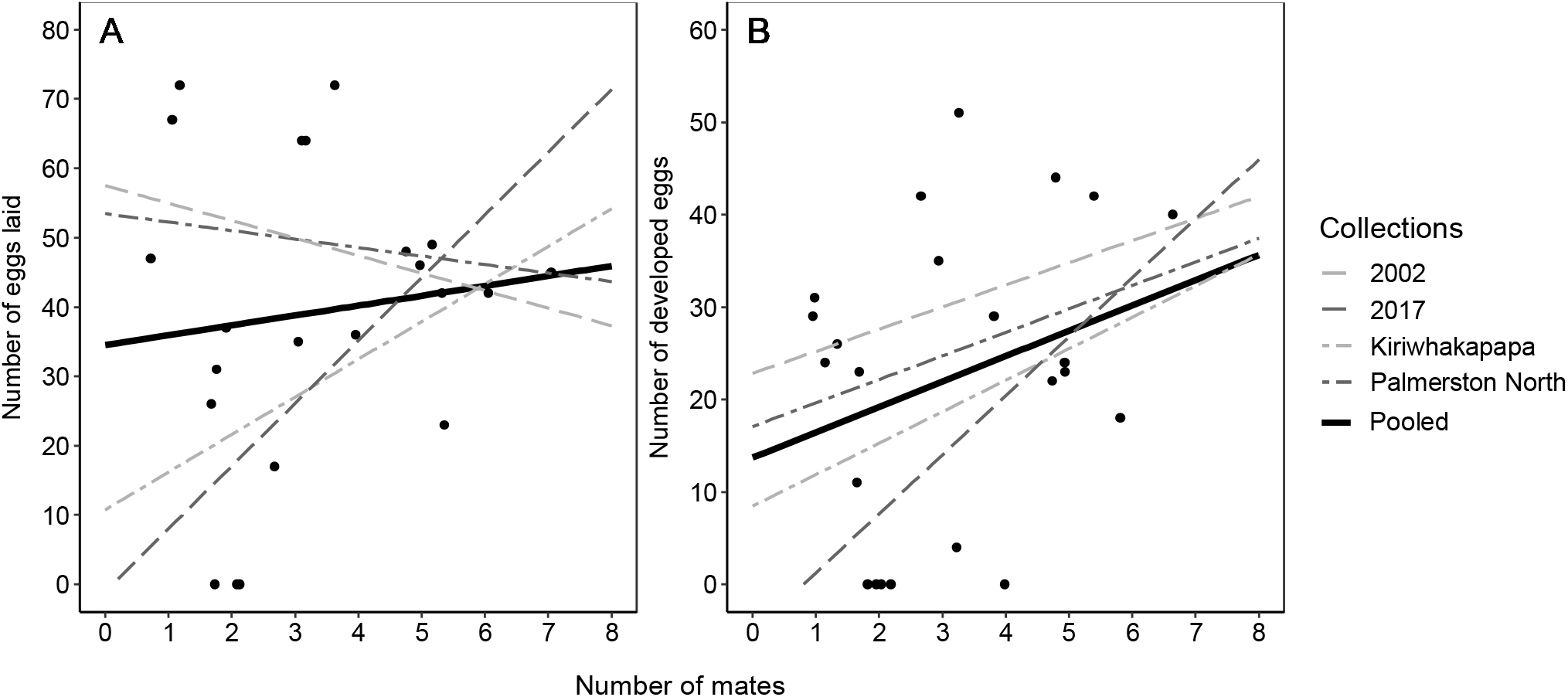
Raw Bateman Gradients for female *H. pallitarsis* showing relationship between number of mates and either A) number of eggs laid or B) number of developed eggs with 95% confidence intervals. Variation between collection sites and years are shown with dashed lines (associated values provided in Online Resource 1: Table S3).

Estimates of β′_ss_ were similar regardless of whether we included the available pronotum data in our regression model (β′_ss_ = 0.155 ± 0.264, F(2, 16) = 1.46, p = 0.280 for eggs laid; β′_ss_ = 0.433 ± 0.310, F(2, 17) = 1.16, p = 0.090 for eggs developed) and in line with Gwynne (2005), female body size had no significant effect on the number of developed eggs (β′ = 2.343 ± 2.982, R^2^ = 0.120, F(2, 17) = 1.16, p = 0.214), but a marginally significant effect on the number of eggs laid (β′ = 4.07 ± 2.490, R^2^ = 0.154, F(2, 16) = 1.46, p = 0.043).

### Sexual selection on accessory organs

Female mating success ranged from 0-6 mates (mean: 2.3 +/- 1.4 SD) and increased significantly with accessory organ length (table 2). Larger body size showed a non-significant trend with increased mating success (table 2). Selection did not vary significantly between collection sites (MLS interaction terms; body size: β′=0.319 ± 0.247, F(3, 50) = 1.39, p=0.202; accessory organ width: β′= -0.331 ± 0.276, F(3,50) = 0.537, p=0.236), but we note that (non-significant) trends tended to vary between our two populations of ground weta (visual provided in Online Resource 1: Fig. S4).

**Table 2.** Relationship between phenotype (traits z-score standardized) and mating success (relative number of mates) in female ground weta, *H. pallitarsis*.

| Female traits | $\beta$ (SE) | p-value |
| --- | --- | --- |
| Body size | 0.106 (0.093) | 0.2094 |
| Accessory organ length | 0.195 (0.097) | 0.0426 * |
| Accessory organ width | -0.023 (0.095) | 0.7776 |

$R^2 = 0.105$ , $F(3, 50) = 1.96$
$\beta$ = standardized directional selection gradients; regression coefficients
SE = Standard error
Estimates obtained from multiple OLS regression; p-values from Permutation tests with 5000 randomizations
\* Statistically significant p-value

None of the accessory organ traits measured significantly predicted the number of eggs in a female’s abdomen. Although we found a statistically significant effect of accessory organ width (table 3), this trend appears to be driven almost entirely by differences over time (visual provided in Online Resource 1: Fig. S5). Indeed, when collection year is added as a random effect in a general linear mixed model, the significant effect of accessory organ width disappears (β′=0.127 ± 0.138, p=0.238). The relationship between female traits and fecundity did not vary significantly between collection sites (OLS interaction terms; body size: β′=0.191 ± 0.342, F(3, 53) = 1.172, p=0.579; accessory organ width: β′= -0.331 ± 0.359, F(3,53) = 2.09, p=0.361, but again (non-significant) trends varied between sites (visual provided in Online Resource: Fig. S6).

**Table 3.**
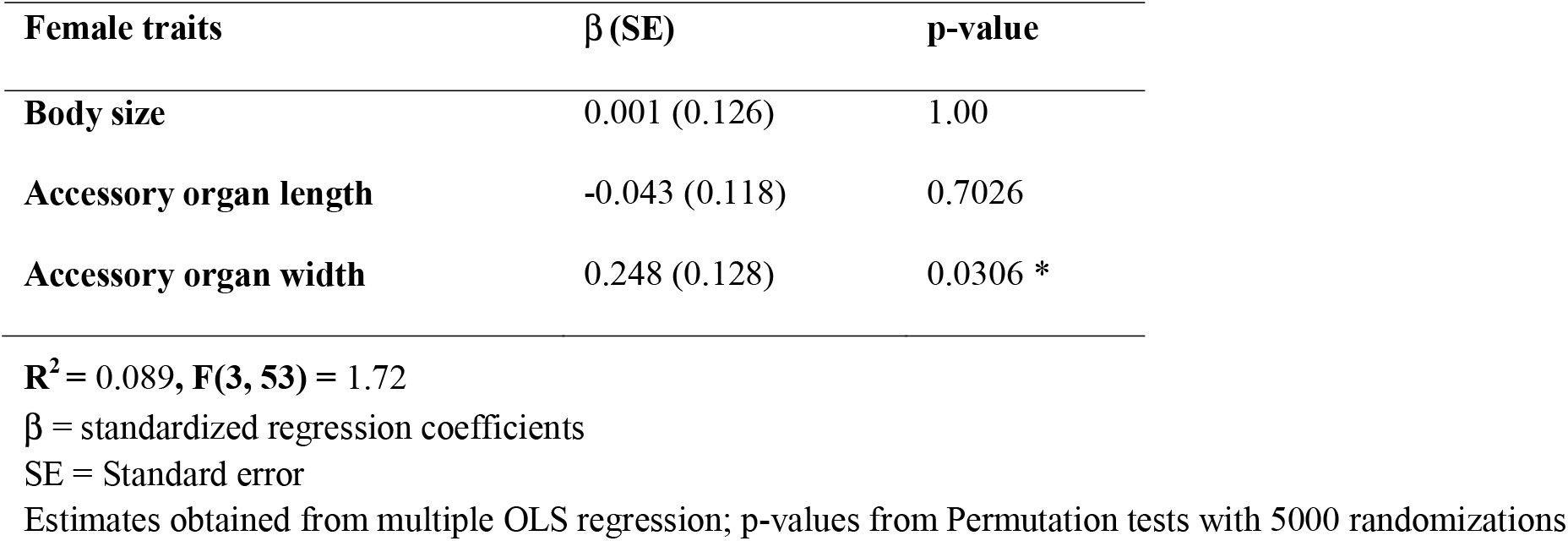
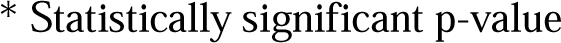
Relationship between female phenotype (traits z-score standardized) and relative number of eggs in the abdomen in female ground weta, *H. pallitarsis*.

| Female traits | $\beta$ (SE) | p-value |
| --- | --- | --- |
| Body size | 0.001 (0.126) | 1.00 |
| Accessory organ length | -0.043 (0.118) | 0.7026 |
| Accessory organ width | 0.248 (0.128) | 0.0306 * |

$R^2 = 0.089$ , $F(3, 53) = 1.72$
$\beta$ = standardized regression coefficients
SE = Standard error
Estimates obtained from multiple OLS regression; p-values from Permutation tests with 5000 randomizations
\* Statistically significant p-value

## Discussion

In *H. pallitarsis* ground weta, sexual competition for male-provided nutrition (spermatophylaxes) prior to an extended maternal care period without food probably creates an opportunity for sexual selection on females when they “forage” for mating gifts. Morphological evidence that females are subject to sexual selection comes from the existence of an apparent female ornament, a secondary-genitalic device known as an accessory organ (Gwynne 2002; Gwynne 2005) that varies greatly between species (Johns 2001; Trewick et al. 2020). Here, we provide some evidence that *H. pallitarsis* females are subject to sexual selection, by showing 1) a significant positive relationship between female mating success and lifetime reproductive success (Standardized Bateman gradient; β′_ss_) and 2) a significant relationship between the length of the female accessory organ and mating success.

Although our estimates of the Bateman gradient were not consistent across collection years, they reached values in 2017 that are similar to other species where sexually selected females compete for mates, such as bean beetles (β′_ss_ =0.62-0.87; Fritzsche & Arnqvist 2013) or pipefish (β′_ss_ = 0.86-1.04; Mobley & Jones 2012; Aronsen et al. 2013). While pooled estimates of the Bateman gradient were not statistically significant, they were higher than the estimates found in female *K. nartee* (β′_ss_ = 0.029-0.034; Hare and Simmons 2022) as well as the relationship between number of sires and absolute reproductive success found for females of another anostostomatid weta, *Hemideina crassidens*, where males are sexually competitive (they fight using very large mandibular weaponry) (β= 0.47; Nason and Kelly 2020).

Further evidence of sexual selection on female ground weta comes from our finding that accessory organs appear to be subject to directional selection. There was a marginally significant positive relationship between the length of the accessory organ and female mating success, supporting the hypothesis that this this structure functions as a device that ensures successful mating (there was also a non-significant trend for larger females to mate more frequently). In all our measures of sexual selection, there was considerable variation between study sites. This is interesting given the lack of genetic differentiation between these populations (F_st_ low compared to typical values in insects; Ward et al 1992) and their close proximity (roughly 100km) relative to the total species range.

Morphological differences have been reported between populations of *H. pallitarsis*, with Chappell et al. (2012) finding that some populations differed in body size as well as the drumming pattern performed by males prior tomating. While this variation was not congruent with genetic differences among populations (suggesting that widely separated *H. pallitarsis* populations comprise one interbreeding species; Chappell et al. 2012), patterns of selection and mate attraction may differ slightly between populations of ground weta in Palmerston North compared to Kiriwhakapapa, possibly because the habitats differ; overgrown gardens (with apparently high population density) in the former site and wild vegetation in the latter. Regardless of this variation, female ground weta appear to be subject to sexual selection, and the length of accessory organ plays a consistent role in acquiring mates in both populations.

Despite this finding, neither accessory organ length, nor any of the female traits measured were related to the number of eggs in a female’s abdomen (fecundity). However, this does not mean the accessory organ does not signal attractiveness to males. One reason for the non-significant finding may be that the number of eggs in the abdomen is not the main indicator of female quality for the mating male. Because females oviposit all their eggs and then engage in a long maternal care period, during which the mother does not eat (Gwynne 2004; Browne and Gwynne 2022), choosy males may prioritise female condition and thus the potential quality of care she is able to provide, as this will determine the number of surviving offspring. This is supported by Gwynne’s (2005) study which showed a positive relationship between accessory organ length and number of hatched nymphs rather than eggs in the abdomen and might also help to explain why our measures of the Bateman gradients tended to be more consistent (and possibly stronger) when using the number of developed eggs to estimate reproductive success rather than just the number of eggs laid (fig. 3; Supplementary information 3). Similar results might also be expected however, if there is variation in quality or size of the spermatophylax gift received, and thus we cannot rule out the possibility that females with longer accessory organs also obtain more nutrients for successful offspring production, due, for example, to male preferences (reviewed in Simmons 2001) or a manipulative anchoring function similar to *Neotroglea* (Yoshizawa et al. 2014; Yoshizawa et al. 2018).

As noted previously, the accessory organ’s role in competing successfully for copulations does not exclude its mechanical function in successful ejaculate transfer (male genitalia facilitate sperm transfer as well as potentially stimulating mates; Eberhard 1990). Given the presumed ancestral state where males used genitalic hooks to latch onto a female’s unmodified 6^th^ sternite, the accessory organ could have initially evolved in the context of mate assessment (in the region that interacts closely with sensitive male genitalia) or as an grasping device between the male genital falci and epiprocts, that facilitates spermatophylax-gift transfer. The association between paired male genitalic structures (falci, paraprocts, and the end of the 8^th^ sternite; D. Gwynne personal observation) almost certainly achieve stability while first the sperm capsule (ampulla) and then the spermatophyax gift is delivered. However, it is possible, that like many male structures (Eberhard 1990), different parts of the accessory organ serve different functions. We note, for example, that the distal end of the accessory organ has a very prominent bilobed shape that appears to align with the lobes of the males’ paired paraprocts (in several *Hemiandrus* species: Gwynne 2002). Because our measures of the accessory organ were simple linear ones, we have not captured the full complexity of this structure that may be important to successful mating. Future studies should focus on quantifying the shape of this structure and its role in mating.

Another limitation of the study is that we were only able to measure directional selection on accessory organs. Due to our limited sample size, we did not compute estimates of non-linear or correlational selection, meaning we may have an incomplete understanding of the type of selection acting on these structures (Lande and Arnold 1983). In particular, it would be valuable to test for stabilizing sexual selection on female accessory organs as preference for intermediately ornamented females has been predicted in some species (typically those with more elaborate and costly traits) due to the potential costs (e.g., in egg production) of investing in these traits (Fitzpatrick 1995; Chenoweth et al. 2006; Wheeler et al. 2012). Furthermore, genital structures may be expected to be under stabilizing sexual selection in cases where they have evolved to fit the average male (Eberhard et al. 1998).

In conclusion, we show some evidence of sexual selection on female ground weta, *H. pallitarsis*, and also on a unique secondary genitalic structure they possess. Directional selection for longer accessory organs suggests this device functions like a sexual ornament to secure matings. Although this structure does not correlate with measures of female fecundity, it may well signal female condition (and thus perhaps her genitalic grasping ability) or quality of care as important predictors of female reproductive success as females engage in a long and costly maternal care period.

## Supporting information

Online Resource 1

## Statements and Declarations

The authors have no competing interests to declare

## Acknowledgements and Funding Statement

We thank Mary Morgan Richards and Lindsay Coome for their help collecting data, Doug Currie, Marc Johnson, Rosalind Murray, and Clint Kelly for comments on the research and manuscript, as well as the New Zealand Department of Conservation (for permits). This research was supported by a Natural Sciences and Engineering Research Council of Canada Discovery Grant to DTG.

