## Supplementary material for "Sexual selection on a female copulatory device in an insect with nuptial gifts": Online Resource 1

Jessica H. Browne^1,2^

Darryl T. Gwynne^1,3^

^1^ Department of Ecology and Evolutionary Biology, University of Toronto Mississauga, Mississauga, Ontario L5L 1C6, Canada

Mount Allison University, 62 York St. Sackville, NB, E4L 1E2, Canada

^3^

**Table S1.** Microsatellite library developed for ground weta *H. pallitarsis* (Genetic Marker Services, Brighton, UK)

| **Locus** | **Primer Sequence (5’-3’)** | **Allele size (bp)** | **Fluro-label** |
| --- | --- | --- | --- |
| **wet14** | F: TTT TCC AAG TGG GTT ATC GTG  R: CGG TAG CTA ATG GTC GGT GT | 169-183 | HEX |
| **wet18** | F: TGC GAT GTT ACC AAA ACA TCA T  R: AGG CGT CCG ATA TCT CCT TT | 96-114 | HEX |
| **wet21** | F: TAG TCC ATT ATT CAA TAC CC  R: GAG AAT AAG AAG TAG CAA AAT AA | 91-127 | 6-FAM |
| **wet22** | F: GTA CTG CAT CTT GTC TGA GAG C  R: AAG AAG TGT CCA AGG CCA TC | 155-235 | 6-FAM |
| **wet27** | F: TCG CAA GAG TAG GCT ACG TG  R: CGT GCG AGT AGG TGT TTG AG | 85-97 | 6-FAM |
| **wet80** | F: TCA ATG AAA TGT TTC AGA CAG ACA  R: GCA CAA AGA GCC ATC TGT CA | 124-172 | HEX |
| **wet81** | F: ACG ACG AGG AAA TGA AAT GT  R: GTT GAC AAA ACT CGG GCA GT | 158-226 | HEX |
| **wet83** | F: GGT AGA CAA AGC GGT TGA TG  R: CAA CAT CCT CCC GTA GTT TT | 120-158 | 6-FAM |
| **wet89** | F: CCC TTG TTC CGT CCT ACA AG  R: CAA AAC CCC ACC ATG ACT TT | 180-258 | 6-FAM |

**Table S2.** Characterization of microsatellite loci for ground weta *H. pallitarsis* collected from two populations

|  | **Kiriwhakapapa** | | | | |  | **Palmerston North** | | | | |  |
| --- | --- | --- | --- | --- | --- | --- | --- | --- | --- | --- | --- | --- |
| **Locus** | **N** | **A** | **H_E_** | **H_O_** | **ƒ_null_** |  | **N** | **A** | **H_E_** | **H_O_** | **ƒ_null_** | **F_st_** |
| **wet14** | 22 | 3 | 0.21 | 0.23 | 0.000 |  | 22 | 5 | 0.58 | 0.77 | 0.019 | 0.16 |
| **wet18** | 17 | 4 | 0.53 | 0.18* | 0.230 |  | 15 | 7 | 0.71 | 0.47* | 0.302 | 0.06 |
| **wet21** | 13 | 9 | 0.80 | 0.31* | 0.303 |  | 15 | 7 | 0.82 | 0.40* | 0.221 | 0.01 |
| **wet22** | 7 | 8 | 0.87 | 0.86 | 0.051 |  | 18 | 13 | 0.88 | 0.83 | 0.000 | 0.04 |
| **wet27** | 19 | 5 | 0.20 | 0.21 | 0.000 |  | 19 | 4 | 0.15 | 0.16 | 0.000 | 0.00 |
| **wet80** | 42 | 17 | 0.91 | 0.93 | 0.002 |  | 31 | 14 | 0.85 | 0.84 | 0.000 | 0.06 |
| **wet81** | 36 | 11 | 0.89 | 0.56* | 0.172 |  | 36 | 20 | 0.92 | 0.92 | 0.007 | 0.03 |
| **wet83bbbbbbbbb** | 31 | 4 | 0.66 | 0.52* | 0.084 |  | 25 | 5 | 0.61 | 0.28* | 0.244 | 0.036 |
| **wet89** | 44 | 16 | 0.90 | 0.77* | 0.085 |  | 31 | 12 | 0.88 | 0.77* | 0.051 | 0.05 |

**N=** number of individuals tested

**A**= number of alleles detected

**H_E_** = Expected heterozygosity

**H_o_** = Observed heterozygosity *significant deviation from Hardy-Weinberg equilibrium

**ƒ_null_ =** estimated frequency of null alleles

All calculations were done using GENEPOP version 4.2 (Raymond and Rousset 1995)

**Table S3.** Bateman Gradients for wild-caught female ground weta, *H. pallitarsi*s shown by collection year and collection site

| **Response variable** | **Sample** | **β′_ss_ (SE)** | **β_ss_ (SE)** | **R2** | **F** | **df** | **p-value** |
| --- | --- | --- | --- | --- | --- | --- | --- |
| **Total eggs laid** | **Pooled** | **0.117 (0.231)** | **1.424 (2.822)** | **0.013** | **0.255** | **1, 20** | **0.619; 0.313** |
|  | 2002 | -0.121 (0.185) | -2.525 (3.839) | 0.067 | 0.432 | 1, 6 | 0.535; 0.249 |
|  | 2017 | 1.032 (0.358) | 9.054 (3.138) | 0.4096 | 8.326 | 1, 12 | 0.0137; 0.0084 * |
|  | Kiriwhakapapa | 0.629 (0.489) | 5.425 (4.215) | 0.155 | 1.656 | 1, 9 | 0.230; 0.114 |
|  | Palmerston North | -0.076 (0.181) | -1.226 (2.902) | 0.019 | 0.178 | 1, 9 | 0.683; 0.330 |
| **Developed eggs** | **Pooled** | **0.392 (0.272)** | **2.745 (1.907)** | **0.090** | **2.071** | **1, 21** | **0.165; 0.084 .** |
|  | 2002 | 0.204 (0.233) | 2.393 (2.739) | 0.098 | 0.764 | 1, 7 | 0.411; 0.205 |
|  | 2017 | 1.277 (0.504) | 6.386 (2.522) | 0.348 | 6.412 | 1, 12 | 0.0263; 0.006 * |
|  | Kiriwhakapapa | 0.575 (0.554) | 3.404 (3.281) | 0.107 | 1.077 | 1, 9 | 0.327; 0.155 |
|  | Palmerston North | 0.315 (0.300) | 2.544 (2.427) | 0.099 | 1.099 | 1, 10 | 0.319; 0.164 |

β′_ss­_ = Standardized Bateman gradient (variables mean-standardized)

β_ss­_ = Raw (unstandardized) Bateman gradient

SE = Standard error

Estimates were obtained from separate univariate OLS regression analyses; p-values derived from OLS and 5000 Permutations are shown, respectively

* Statistically significant p-value

Pooled

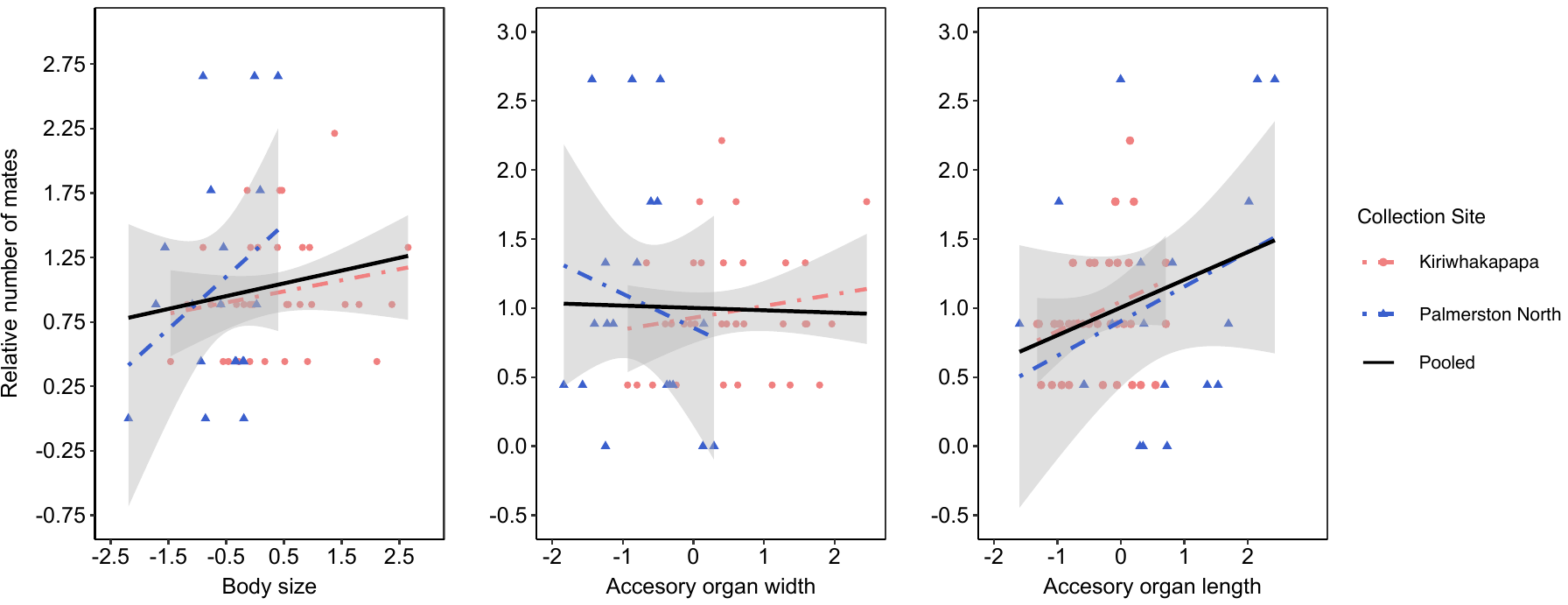

**Figure S4.** Relationship between female (*H. pallitarsis*) traits (z-score standardized) and relative mating success, shown by collection site with 95% confidence intervals, and pooled

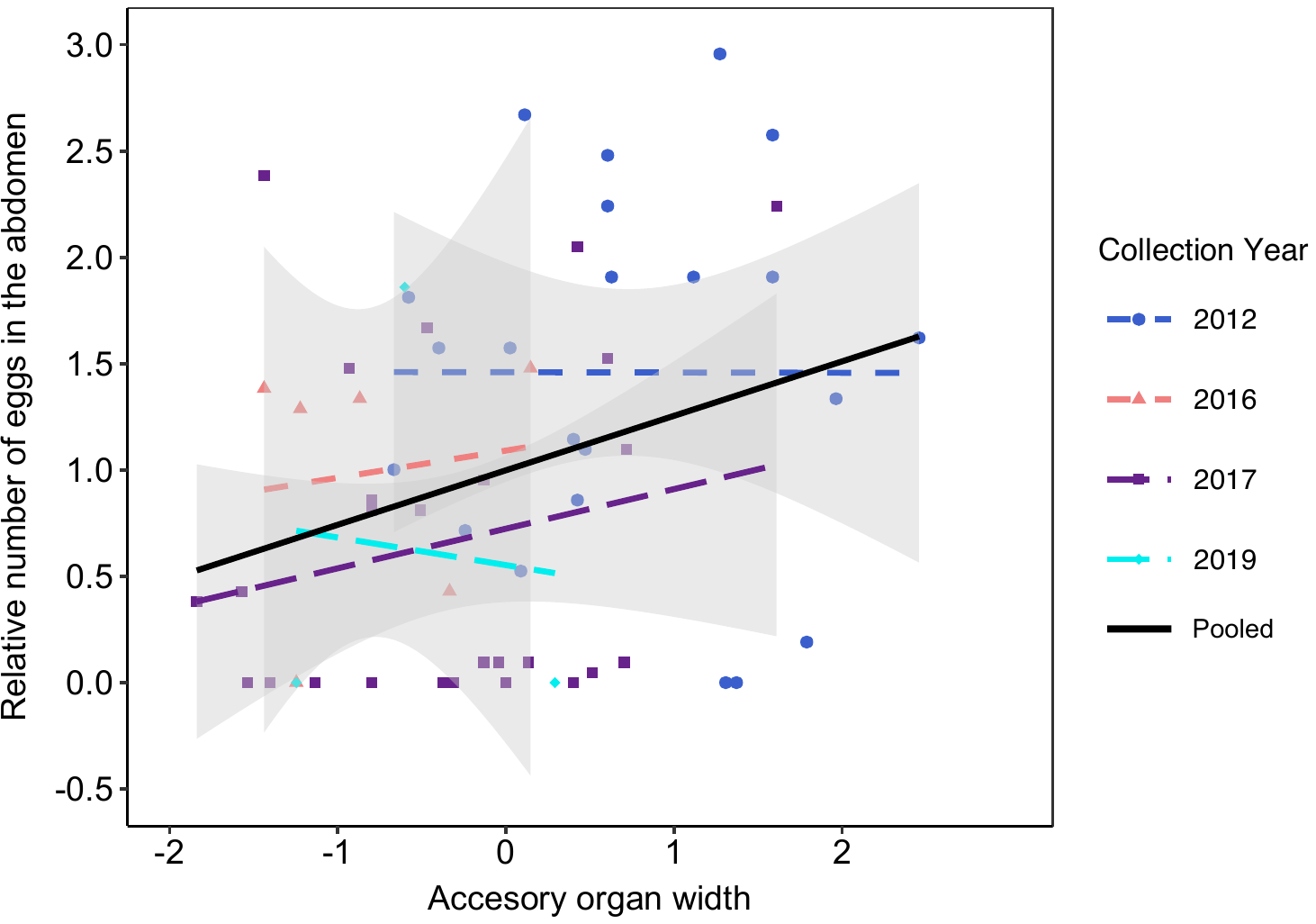

Pooled

**Figure S5.** Relationship between female accessory organ width (z-score standardized) and relative fecundity (number of eggs in the abdomen), shown by collection year with 95% confidence intervals, and pooled

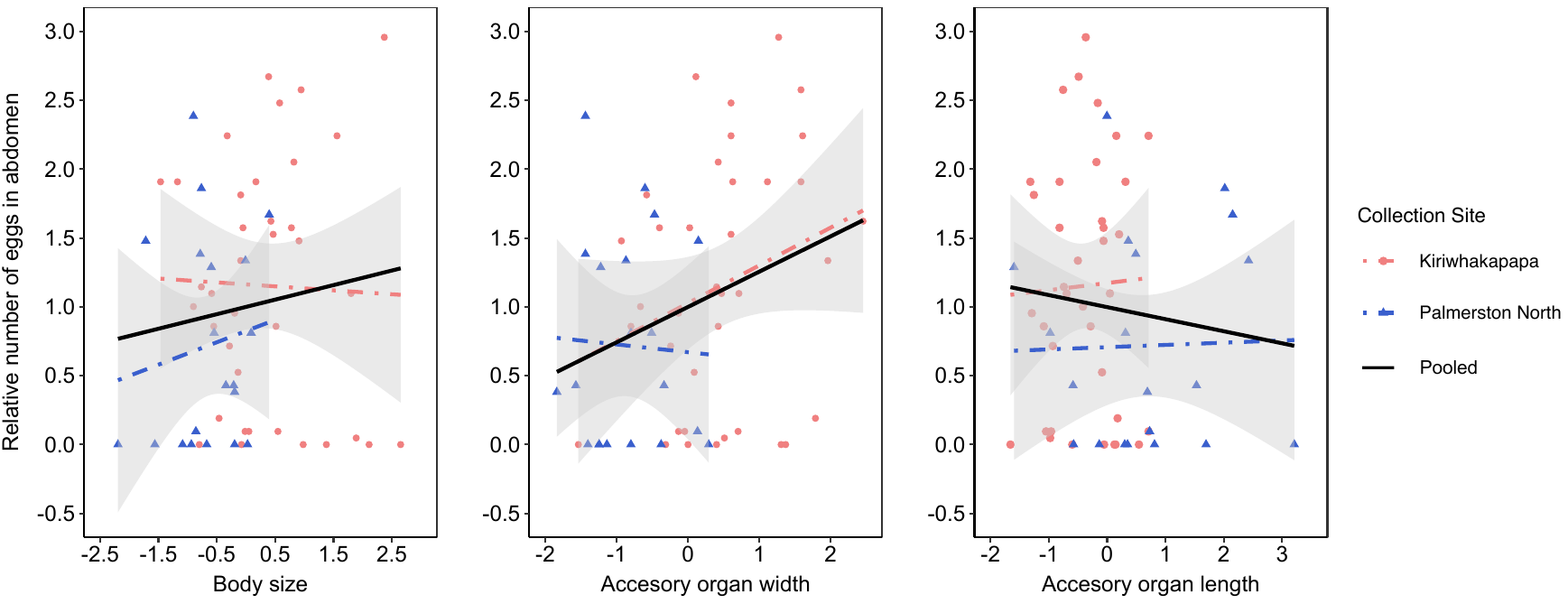

**Figure S6.** Relationship between female (*H. pallitarsis*) traits (z-score standardized) and relative fecundity, shown by collection site with 95% confidence intervals, and pooled
